# MAP7 recruits kinesin-1 to microtubules to direct organelle transport

**DOI:** 10.1101/557611

**Authors:** Abdullah R. Chaudhary, Hailong Lu, Elena B. Krementsova, Carol S. Bookwalter, Kathleen M. Trybus, Adam G. Hendricks

## Abstract

Microtubule-associated proteins (MAPs) play well-characterized roles in regulating microtubule polymerization, dynamics, and organization. In addition, MAPs control transport along microtubules by regulating the motility of kinesin and dynein. MAP7 (ensconsin, E-MAP-115) is a ubiquitous MAP that organizes the microtubule cytoskeleton in mitosis and neuronal branching. MAP7 also promotes the interaction of kinesin-1 with microtubules. We expressed and purified full-length kinesin-1 and MAP7 in Sf9 cells. In single-molecule motiity assays, MAP7 recruits kinesin-1 to microtubules, increasing the frequency of both diffusive and processive runs. Optical trapping assays on beads transported by single and teams of kinesin-1 motors indicate that MAP7 increases the relative binding rate of kinesin-1 and the number of motors simultaneously engaged in ensembles. To examine the role of MAP7 in regulating bidirectional transport, we isolated late phagosomes along with their native set of kinesin-1, kinesin-2, and dynein motors. Bidirectional cargoes exhibit a clear shift towards plus-end directed motility on MAP7-decorated microtubules due to increased forces exerted by kinesin teams. Collectively, our results indicate that MAP7 enhances kinesin-1 recruitment to microtubules and targets organelle transport to the plus end.

## Introduction

Intracellular cargoes navigate a complex network of microtubule tracks to find their destination in the cell. Many cargoes (e.g. endosomes, mitochondria, peroxisomes, mRNA) are transported by similar sets of kinesin and dynein motors, yet they exhibit different transport characteristics and localizations. What cues guide a specific cargo to its destination in the cell? Mounting evidence suggests a role for the microtubule cytoskeleton in directing intracellular transport (1) through the organization of the microtubule network (2–4), tubulin post-translational modifications (PTMs) (2, 5–7), and microtubule-associated proteins (MAPs).

MAPs regulate microtubule dynamics and organization (8–10). MAPs also control the interface between motor proteins and the microtubule surface to regulate motility. For example, the neuronal MAP tau reduces kinesin-1 processivity (11, 12). On cargoes transported bidirectionally by opposing kinesin and dynein motors, inhibition of kinesin-1 biases transport towards the microtubule minus end (13). MAP4 inhibits kinesin-1 motility in gliding assays (14) and regulates the aggregation and dispersion of pigment granules transported by kinesin-2 and dynein (15). Doublecortin and doublecortin-like kinase 1 promote the motility of kinesin-3 motors in neurons (16, 17). Thus, MAPs can exert tight control on transport as their effects are specific to each MAP and motor protein.

Here, we focus on the role of MAP7 (ensconsin, E-MAP-115) in regulating intracellular transport. MAP7 is a microtubule-associated protein involved in a diverse array of cellular processes. By serving as a microtubule polymerase (18, 19), MAP7 enhances neuronal branching and controls the spindle length (20). MAP7 strongly affects kinesin-1 motility. Loss of MAP7 in oocytes resulted in impaired plus-ended motility (21). In single-molecule motility assays, MAP7 increases the frequency of processive runs by kinesin-1 (21–24) but does not affect dynein motility (23).

MAP7’s strong effects on kinesin-1 motiity suggest a role in regulating the transport of intracellular cargoes transported by kinesin and dynein motors. We used single-molecule motility assays and optical trapping to measure the effect of MAP7 on the motility of single full-length kinesin-1, teams of kinesin-1, and native sets of kinesin and dynein motors on isolated organelles. Optical trapping assays indicate that MAP7 increases the binding rate of kinesin-1 to microtubules, but does not affect the force generation or processivity of single kinesin motors. For teams of kinesin-1, this increased binding rate results in a greater number of motors being engaged with the microtubule at any given time, resulting in enhanced processivity and force generation. On native cargoes transported by teams of kinesin-1, kinesin-2, and dynein, kinesin motors produce higher forces in the presence of MAP7 to target transport towards the microtubule plus end.

## Results

### MAP7 directs bidirectional cargoes to the microtubule plus end

Given MAP7’s role as an activator of kinesin-1, we hypothesized that it might target the motility of bidirectional cargoes towards the microtubule plus end. We isolated phagosomes along with their native transport machinery, and reconstituted their motility along polarity-marked microtubules. Phagosomes are transported by teams of 1-3 kinesin-1, 2-6 kinesin-2, and 3-10 dynein motors (13, 25).

Using an optical trap, we measured the forces exerted by teams of kinesin and dynein motors. In the absence of MAP7, isolated phagosomes spend an approximately equal fraction of time moving towards the microtubule plus or minus end (Fig. 1C,D) (13, 26). MAP7 increases the frequency and magnitude of kinesin-directed force events, while the frequency and magnitude of dynein-directed forces decrease. (Fig. 1C, S1C,E).

**Fig 1.**
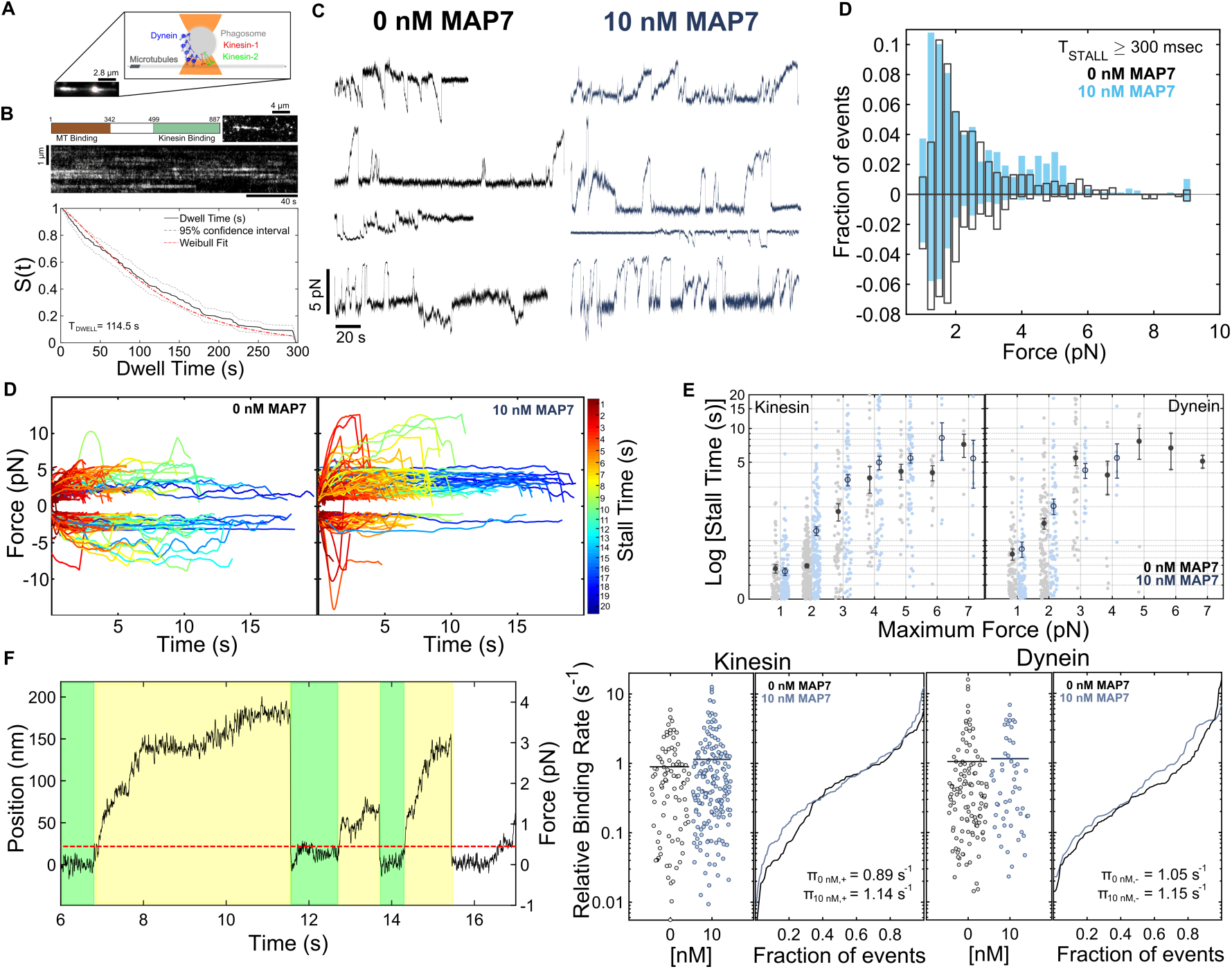
MAP7 targets bidirectional cargoes to the microtubule plus end by increasing the number of engaged kinesin motors. **(A)** Isolated phagosomes containing fluorescent beads were positioned on polarity-marked microtubules and the forces were measured using an optical trap. These isolated phagosomes are transported by teams of kinesin–1, kinesin–2, and dynein motors (13). **(B)** MAP7 forms static patches along the microtubule lattice. Using survival analysis, we determined the average dwell time of MAP7 on microtubules. Mean dwell time of MAP7 on microtubules is 114.5 +/-6.16s (*n* = 221 events from 11 recordings, 2 independent experiments). **(C)** Force traces were acquired at 2kHz and median filtered at 20 Hz. Maximum forces for all trap displacements greater than 300 ms in duration were recorded (0 nM: *n*=689 events from 38 recordings, 11 independent experiments; 10 nM MAP7: *n*=787 events from 40 recordings, 11 independent experiments). Consistent with previous results (13, 26), plus–end directed force events consist of unitary stall forces of kinesin–1 and kinesin–2, events where the motors detach before reaching their stall force, and rare events driven by multiple kinesins. Minus–end directed forces indicate events driven by teams of several dynein motors (Fig. S1C). The frequency and magnitude of kinesin-driven forces are increased in the presence of 10 nM MAP7. In response, dynein-mediated forces are reduced. The Bayesian Information Criterion was used to determine the optimal number of components to describe the force histograms (Fig. S1F). Mean forces of the multicomponent fits for plus–end directed forces are 1.57, 2.38, 4.5, and for minus-end directed forces are 1.14, 1.6, 2.8, 5.8 pN. With MAP7, mean forces of the multicomponent fits for plus-end directed forces are 1.4, 2.4, 4.5, 7.3 pN and for minus-end directed forces are 1.4, 2.7, ≥9 pN. **(D,E)** Teams of kinesin and dynein motors remain attached to the microtubule under load for longer durations in the presence of MAP7. The color of the trajectory indicates the duration of a stall event with short events color coded by red and long events color coded by blue. The stall time is significantly higher for kinesin motors, when compared to dynein motors on isolated phagosomes, indicating a longer engagement time of the microtubules (*< T*_0*nM,*+_ *>*=1.5s, *< T*_10*nM,*+_ *>*=2.5s (*p<*0.01); *< T*_0*nM,-*_ *>* = 2.26s, *< T*_10*nM,-*_ *>* = 1.8s (*p*=n.s.). Statistical test = Student’s t-test). **(F)** The relative binding rate was calculated as the reciprocal of the dwell time preceding a force event. The green area indicates periods where the phagosome is diffusing in the optical trap and yellow areas indicate processive force events (Appendix S2). MAP7 increases the relative binding rate of plus-ended motors on MAP7 decorated microtubules by 30%.

We next examined the processivity of teams of kinesin and dynein motors as a function of opposing load. We observe that cargoes driven along MAP7-decorated microtubules stay engaged for longer periods of time before detaching from the microtubules. The duration plus-ended motors remain engaged before detaching (stall time) increases by ∼ 67%, while the average stall time of minus-ended motors decreases only by ∼ 18% (Fig. 1D). We compared motility by similar numbers of motors by examining the duration of force events with similar magnitudes, and conclude that MAP7 does not significantly affect the processivity of individual kinesin or dynein motors (Fig. 1E). We calculated the force-dependent unbinding rate using the method proposed by Berger et al. (2018), and find the unbinding rate for both kinesin and dynein is unaffected by MAP7 (Fig. S1G). These analyses indicate that while MAP7 does not alter the processivity of individual motors, it does increase the processivity of kinesin teams by increasing the number of engaged motors.

To compare the effect of MAP7 on the rate at which kinesin and dynein motor teams bind to microtubules under similar conditions, we estimated the relative binding rate from the duration of diffusive dwell preceding motor-driven events in the optical trap. MAP7 increases the relative binding rate of plus-ended motors by ∼ 30%, while the binding rate of dynein teams is increased by 10% (Fig. 1F, Appendix 2). Additionally, the number binding events per recording increases for kinesin-1 motors and decreases for dynein motors in the presence of MAP7 (Fig. S1D). Taken together, these results suggest that MAP7 enhances kinesin binding, such that a larger fraction of the kinesin motors are engaged at any time and thus exert greater forces.

### MAP7 recruits kinesin-1 to microtubules

To dissect the mechanism through which MAP7 regulates the activity of plus-ended motors, we performed single-molecule motility assays with full length kinesin-1 (Fig. 2A). MAP7 increases the binding rate of kinesin-1 motors on the microtubules in a dose dependent manner (Fig. 2B). While kinesin-1 run lengths increase (Fig. 2C), their velocities (Fig. S2A) are largely unchanged in the presence of MAP7. MAP7 binds to kinesin-1 on the stalk domain, near the hinge region that mediates auto-inhibition (19, 23, 28). Thus, we asked if MAP7 acts both to recruit kinesin-1 to microtubules and relieve auto-inhibition (29, 30). In single-molecule motility assays with full-length kinesin-1, MAP7 increases both the number of processive and diffusive runs to a similar degree, suggesting that MAP7 does not regulate kinesin autoinhibition. The fraction of diffusive runs increases at high levels of MAP7 concentration (75 nM), indicating that MAP7 can act as an obstacle on the microtubule when present at high densities (Fig. 2D, S2B). Motility assays using *Drosophilia* kinesin-1 and MAP7 (Fig. 2A-D) or mammalian kinesin-1 and MAP7 (Fig. S2C-H) show similar results.

**Fig 2.**
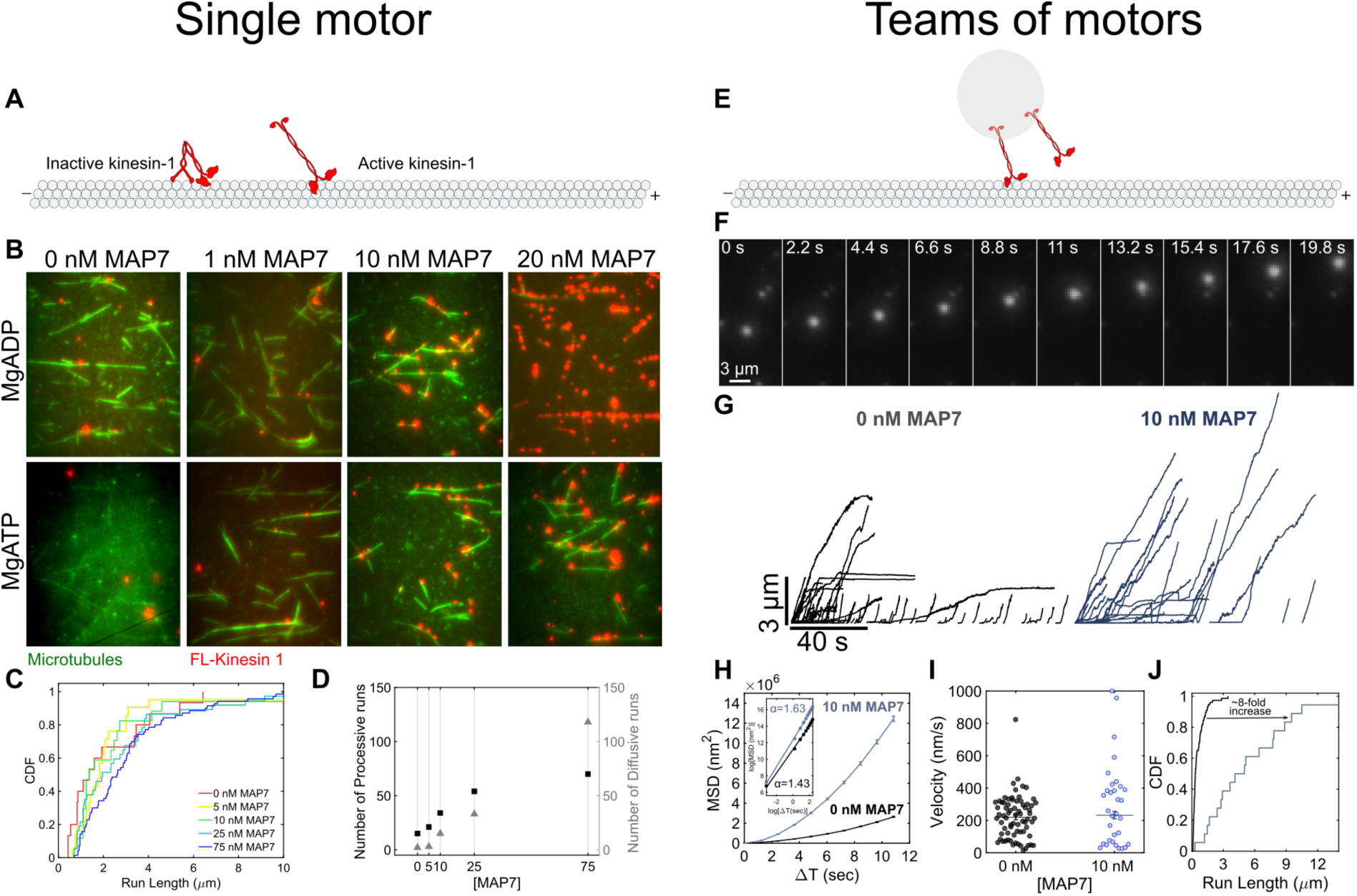
MAP7 recruits kinesin-1 to microtubules. **(A,B)** MAP7 recruits full-length kinesin–1 to microtubules in a dose–dependent manner (Fig. S2B). The motility of single molecules of kinesin-1 was reconstituted along MAP7-decorated microtubules. Due to auto-inhibition, full-length kinesin-1 exhibits both diffusion and processive runs. **(C)** At high levels of MAP7, kinesin-1 run lengths increase by ∼ 30%, possibly due to rapid re-attachment (Average run length: 0 nM = 2033 +/-493.8 nm; 5 nM = 1640 +/-463 nm; 10 nM = 1800 +/-937 nm; 25 nM = 2227 +/-411.2 nm; 75 nM = 3002 +/-265.5 nm). **(D)** The frequency of both processive and diffusive motility of kinesin-1 increases to a similar degree with increasing MAP7 concentration. (0 nM MAP7: *n*= 15 runs 3 recordings; 5 nM MAP7: *n*= 17 events from 3 recordings; 10 nM MAP7: *n*= 21 events from 3 recordings; 25 nM MAP7: *n*= 37 events from 3 recordings; 75 nM MAP7: *n*= 70 events from 3 recordings). **(E,F)** We next attached kinesin–1 to anti-GFP fluorescent beads. Motor-coated beads were positioned on the microtubule via optical trapping and were allowed to escape from a weak optical trap (k∼ 0.004 pN/nm). **(G)** Long directed runs towards the microtubule plus-end indicate motility by multiple kinesin–1 motors. **(H)** Teams of kinesin–1 motors are more processive on MAP7 decorated microtubule **(I,J)**. The average run length of kinesin–1 motors with MAP7 increases by ∼8 fold (Average run length (0 nM) = 595 +/-60 nm; Average run length (10 nM) = 5320 +/-890 nm), while the velocity is unaffected (Average velocity (0 nM) = 220 +/-16 nm/s; Average velocity (10 nM) = 292 +/-45 nm/s). (0 nM: *n*=72 runs, 21 trajectories, 2 independent experiments; 10 nM MAP7: *n*=32 runs from 18 trajectories, 2 independent experiments).

We next asked how enhanced recruitment by MAP7 might affect transport by teams of kinesin-1 motors. We prepared latex beads bound to mammalian full length kinesin-1 motors and titrated the kinesin-1:bead concentration such that we observe ∼ 1-2 active kinesin-1 motors (Fig. 2E,F; *Methods*). The average run length for teams of ∼ 1-2 kinesin-1 motors is 594 nm (Fig. 2G), similar to the run length of a single kinesin-1 motor (31, 32). With MAP7, teams of kinesin-1 motors are more processive as indicated by MSD analysis (Fig. 2G,H) and there is an 8-fold increase in run length (Fig. 2J). Our results, along with previous studies (21–23), suggest that MAP7 enhances the recruitment of kinesin-1 to microtubules. Furthermore, for cargoes transported by multiple kinesins, MAP7 increases the number of motors engaged to increase cargo processivity.

### Teams of kinesin-1 motors exert greater forces on MAP7-decorated microtubules

To investigate how mod-ulating the binding rate of kinesin-1 translates to changes in motility by teams of motors, we performed optical trapping on beads transported by single and teams of mammalian full length kinesin-1 motors (Fig. 3A,G,M, S3S). We tested three conditions: 5000 kinesin-1/bead (single engaged kinesin-1), 10000 kinesin-1/bead (∼ 1-2 engaged kinesin-1s), and 15000 kinesin-1/bead (∼ 2-3 engaged kinesin-1s).

**Fig 3.**
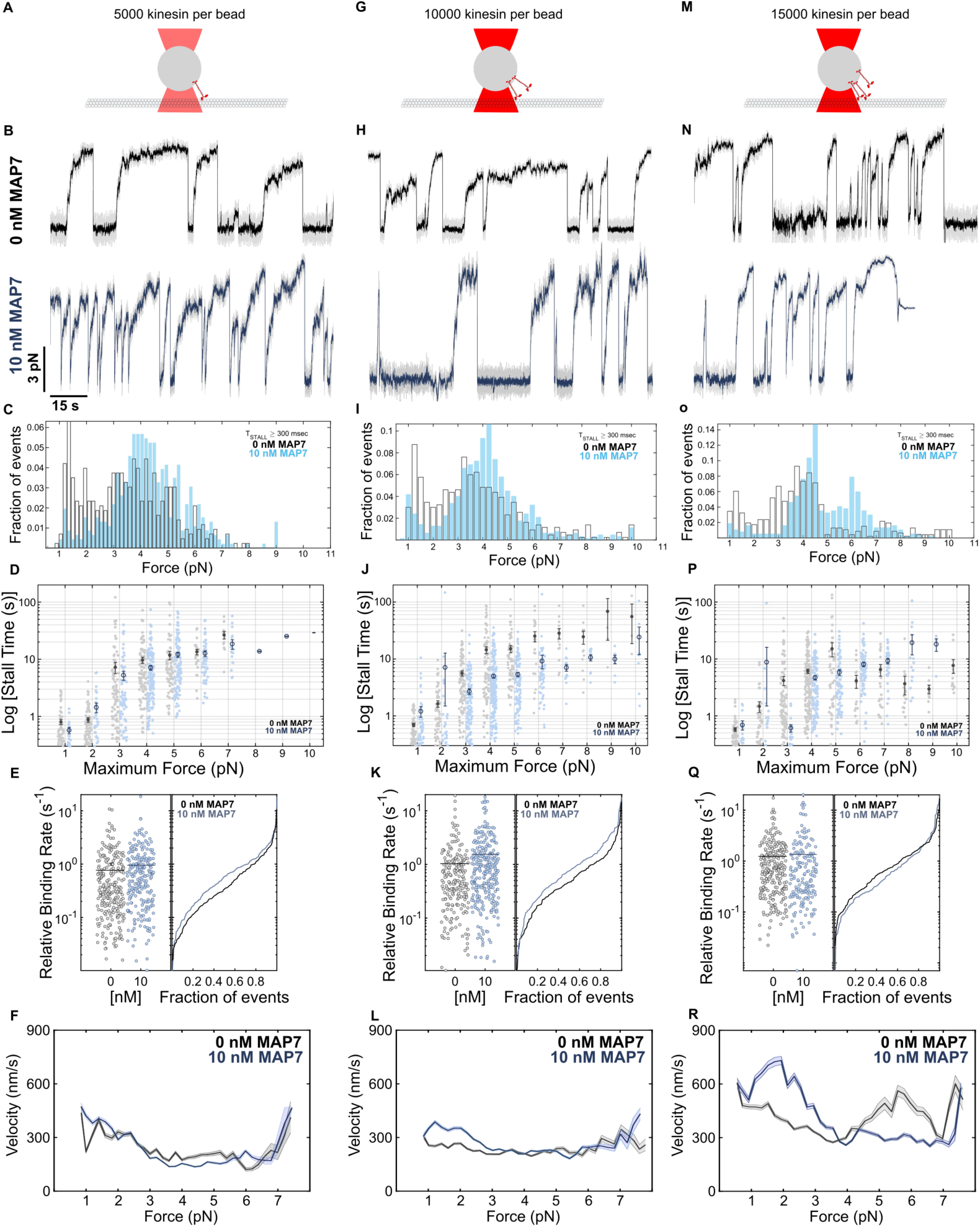
MAP7 increases the forces generated by teams of kinesin-1. We measured forces by single and teams of kinesin–1 motors at three motor densities. **(A)** 5000 kinesin–1*/*bead resulted in motility driven by a single kinesin–1, where *<* 50% of beads interacted with the microtubule; **(G)** 10,000 kinesin–1*/*bead resulted in motility due to 1–2 engaged kinesin–1 motors; **(M)**, and 15000 kinesin–1*/*bead resulted in 2–3 kinesin-1 motors. **(B,C)** On beads driven by single kinesin–1 motors, force distributions indicate unitary stall force of kinesin-1 and sub-stall detachment forces (Mean forces of the multicomponent fits *F*_*comp*_=1.3, 4.03 pN). With the addition of MAP7, sub-stall detachment events decrease and the frequency of unitary stall force events due to single kinesin-1 increases (*F*_*comp*_ = 4.23, *≥* 9pN). (0 nM: *n*=444 events from 25 recordings, 4 independent experiments; 10 nM MAP7: *n*=536 events from 25 recordings, 4 independent experiments). **(H,I)** On beads driven by approximately 1-2 kinesin-1s, force distributions show three distinct populations - forces at unitary stall force of kinesin-1, detachment forces, and rare multimotor events (*F*_*comp*_= 1.27, 3.47, 7 pN). With the addition of MAP7, detachment and low force events decrease, while the frequency of unitary stall force events increase (*F*_*comp*_=1.17, 4, 7 pN). The frequency and magnitude of multimotor force events remain unchanged (0 nM: *n*=479 events from 26 recordings, 4 independent experiments; 10 nM MAP7: *n*=623 events from 24 recordings, 4 independent experiments). **(N,O)** On beads driven by ∼2-3 kinesin-1 motors, force distribution shows three distinct populations - forces at unitary stall force of kinesin-1, low-force events, and multimotor events (*F*_*comp*_= 1.24, 3.64, 7.5 pN). With MAP7, we observe a shift towards frequent high-force events driven by multiple engaged kinesin-1s (*F*_*comp*_= 4.24, 5.97, 7.29) (0 nM: *n*=485 events from 23 recordings, 3 independent experiments; 10 nM MAP7: *n*=378 events from 22 recordings, 3 independent experiments). **(D,J,P,S3)** The duration of stall events at a given force is not strongly influenced by MAP7, indicating that MAP7 does not affect the processivity of single kinesin motors. **(E)** Single kinesin-1 motors and **(K)** teams of 1-2 kinesin-1 motors bind much faster to MAP7-decorated microtubules. **(Q)** The increase in binding rate is not observed for larger teams of 2-3 kinesin-1, likely because attachment is no longer limited by the single motor binding rate when many motors are available for binding (Single kinesin-1: 0 nM=0.77*s*^-1^ and 10 nM=0.95*s*^-1^; Teams of 1-2 kinesin-1: 0 nM=1.04*s*^-1^ and 10 nM=1.55*s*^-1^; ∼2-3 kinesin-1: 0 nM=1.24*s*^-1^ and 10 nM=1.35*s*^-1^). **(F,L,R)** The force-velocity curves indicate that more kinesin-1 motors are simultaneously engaged when MAP7 is present, as indicated by higher velocities at the same load.

Force histograms for single kinesin-1 motors show two populations: low-force (∼ 1-2 pN) events due to motor detachment and stall events (∼ 5 pN). In the presence of MAP7, the frequency of low force events decreases and stall force events are more frequent (Fig. 3B,C). Similar results were obtained for beads transported by 1-2 motors, with the exception of rare multimotor events (Fig. 3H,I). For teams of 2-3 kinesin-1 motors, we observe 3 groups of force events - low force detachment events, stall force events, and high-force events due to multiple motors (Fig. 3N,O). In the presence of MAP7, low-force detachment events are rare, and stall force events shift to higher forces, indicating more motors are engaged (Fig. 3O).

In agreement with the results obtained for isolated phago-somes, MAP7 enhances kinesin-1 activity by increasing the binding rate. Attachment time for single (Fig. 3D, S3F) and teams of ∼ 2-3 kinesin-1 (Fig. 3P, S3R) remains unaffected in the absence or presence of MAP7. On the other hand, teams of ∼ 1-2 kinesin-1 remain attached for a shorter interval for a given force (Fig. 3J, S3L). To explain this behavior, we looked at the relative binding rates. The relative binding rate increases by ∼ 25% for single kinesin-1 motors in the presence of MAP7 (Fig. 3E), and ∼ 49% for teams of 1-2 kinesin-1 motors (Fig. 3K). This increase in the binding rate of kinesin motors on isolated phagosomes is small compared to kinesin-1 coated beads, likely due to the presence of kinesin-2 motors, whose binding to the microtubules is unlikely to be affected by MAP7 (Fig. S4). Consistent with the endogenous cargo data, optical trapping data from single and teams of ∼ 1-2 kinesin-1 demonstrate an observable increase in the kinesin-1 binding rate. When many motors are present, the binding rate is saturated and not limited by the binding rate for a single motor. As evidence, the binding rate for ∼ 2-3 kinesin-1 motors in the absence of MAP7 is 1.24 s^-1^ and in the presence of MAP7 is 1.35 s^-1^ (Fig. 3Q). These results suggest that kinesin-1 binding to microtubules is regulated by MAPs. Processivity, as determined by the number of motors engaged to a microtubule and the amount of force they exert, also increases for cargoes driven by single and teams of similar and opposite polarity motors.

Teams of kinesin-1 motors do not share loads equally (32, 33); leading kinesin-1s carry a greater proportion of the load than trailing kinesins. To analyze how MAP7 affects this load sharing between teams of kinesin-1 motors, we looked at their force-velocity relation (Appendix 4, Fig. S3T). Cargoes transported by kinesin-1 teams move faster at a given load in the presence of MAP7, indicating that by increasing the number of active kinesin-1 bound to the microtubule, MAP7 enables loads to be more equally distributed among kinesin-1 teams and increases their ability to carry opposing load (Fig. 3F,L,R). Note that in the case where cargoes are driven by ∼2-3 kinesin-1 motors, the velocity is greater for the control case in the range of loads ∼5-6 pN, likely because high forces are only observed when multiple motors are engaged, while the high force events in the MAP7 case are due to a combination of events driven by single or multiple engaged kinesins. Collectively, these results demonstrate that MAP7 recruits kinesin-1 to the microtubule and a greater number of engaged motors enables kinesin teams to exert higher forces (Fig. 3C,I,O) and move more processively (Fig. 2G,H,J).

### MAP7 regulates bidirectional transport by tuning the kinesin-1 binding rate

Bidirectional motility has been modeled as a stochastic tug-of-war, where force-dependent unbinding of opposing motor teams results in directional switches and pauses (34). We previously extended this model to describe the interactions between teams of three motors - dynein, kinesin-1, and kinesin-2 (13). To quantitatively relate our observations of the effect of MAP7 on single kinesin motors and organelles driven by teams of kinesin and dynein motors, we applied the increase in the binding rate of single kinesin-1 motors we observed experimentally (Fig. 3E) to a simulation of transport by 1 kinesin-1, 2 kinesin-2, and 10 dynein motors (Fig. 4A). Dynein motility is largely unaffected by MAP7 (23). To assess the possibility of an interaction between MAP7 and kinesin-2, we performed a global alignment of kinesin-1 (Heavy Chain) and kinesin-2 motors as MAP7 interacts with the stalk domain of the kinesin-1 heavy chain (Fig. S4A) (19). We observe no homology in the kinesin-1 and kinesin-2 stalk domains, indicating that MAP7 is unlikely to directly interact with kinesin-2. When the enhanced binding rate for kinesin-1 by MAP7 is applied to the model of phagosome transport, motility dramatically shifts towards the microtubule plus-end (Fig. 4B,D). The length of processive runs of kinesin teams increases while the processivity of dynein teams decreases (Fig. 4C). Next, we quantified the number of active motors and their probability distribution in the simulations, *p*(*n*_*kinesin*_,*n*_*dynein*_). In the absence of MAP7, we observe that kinesin and dynein motors are approximately equally likely to be engaged. With MAP7, the probability of kinesin-1 teams being engaged increases while dynein teams are engaged less often (Fig. 4E).

**Fig 4.**
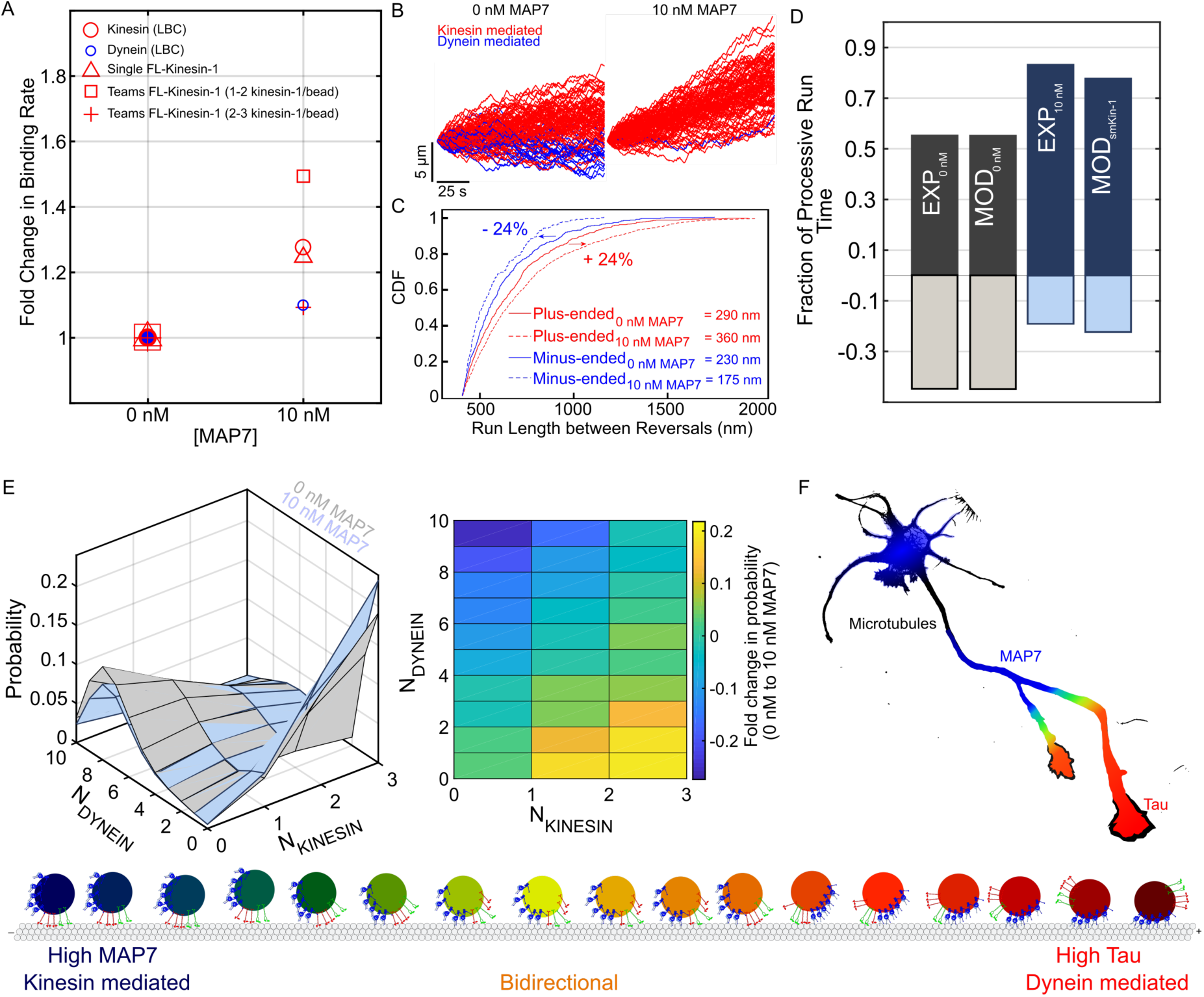
MAP7 directs transport towards the microtubule plus-end. (A) We extended the mathematical model proposed by Muller et al. (34) to describe the interaction between teams of kinesin-1, kinesin-2, and dynein motors based on single motor parameters including the motor stall force, detachment force, unbinding and binding rates (13). We modelled the effect of MAP7 by increasing the binding rate of single kinesin-1 molecules to the same degree as was measured in our optical trapping measurements. Kinesin-2 and dynein were assumed to be unaffected by MAP7. **(B)** The increase in kinesin-1 binding rate biases simulated trajectories towards the plus-end (Number of simulated trajectories, *n* = 100). **(C)** We calculated run length as distance between reversals (25) and consider runs *>* 400 nm as processive. Plus-end directed run lengths increase by ∼24% while minus-end directed run lengths decrease by ∼24%. **(D)** The model results suggest that the shift in phagosome transport towards the microtubule plus end that we observe can be described by simply increasing the kinesin-1 binding rate with no direct effect on kinesin-2 or dynein (EXP = experiment, MOD = model). **(E)** The number of engaged kinesin and dynein motors changes in the presence of MAP7. A greater number of plus-ended motors are simultaneously engaged and exerting force, and as a result minus-ended motors are under higher loads and are less processive. **(F)** MAP7 and tau exhibit distinct localizations in neurons. While MAP7 is enriched at axonal branches (19), tau is localized in a gradient along the axon. Thus, MAP7 might target cargoes to the microtubule plus-end at branch sites, whereas tau directs distal cargoes towards the cell body.

## Discussion

MAP7 promotes binding of kinesin-1 to microtubules to enable a range of kinesin-1 mediated roles, including intra-cellular transport, oocyte polarization, nuclear positioning, and centrosome separation (18, 19, 21–24, 35). Given its central role in regulating kinesin-1 activity, we sought to uncover the mechanisms through which MAP7 controls organelle transport. We quantified the effect of MAP7 on the motility and force generation of single molecules of full length kinesin-1, teams of kinesin-1, and organelles transported by kinesin-1, kinesin-2, and dynein motors. Our results suggest that MAP7 accelerates kinesin-1 binding to microtubules. On cargoes driven by single kinesin-1 motors, MAP7 increases the binding rate but does not alter processivity or force generation. For teams of kinesin-1, the increased binding rate results in a greater number of the motors associated with the cargo being engaged with the microtubule. This leads to enhanced processivity and higher forces exerted by teams of kinesin-1 motors. Vesicular cargoes and organelles are often transported by teams of opposing kinesin and dynein motors. Isolated phagosomes, driven by teams of kinesin and dynein motors, move with approximately equal fraction of plus-and minus-end directed motility. On MAP7 decorated microtubules, kinesin teams produce higher forces such that ∼80% of motility is directed towards the microtubule plus end.

In vitro motility assays (32, 36) and optical trapping in living cells (26, 37) indicate that even when multiple kinesins are bound to a cargo, only a single motor is often engaged with the microtubule at a time. In agreement, we observe that teams of motors rarely exert forces *>* 5 pN (Fig. 3C,I,O). However, when MAP7 is present, the frequency of high-force events increases and that of low-force events decreases. These results suggest that multiple kinesin-1 motors on a single cargo are able to engage simultaneously (Fig. S3B,C) and negative interference between trailing and leading kinesin-1 motors can be overcome by increasing the binding rate. As a result, MAP7 reduces the effect of unequal load sharing among kinesin-1 teams, thus increasing their ability to share opposing load (Fig. 3F,L,R). Mathematical modeling suggests that the enhanced binding rate we observe for a single kinesin-1 motor (Fig. 4E) results in the ability of more motors to engage when operating as a team.

Previous studies have demonstrated that MAP7 is an essential regulator of kinesin-1 dependent transport in vivo (21, 22). We reconstituted the motility of isolated phagosomes driven by teams of kinesin-1, kinesin-2, and dynein motors. Phagosomes are transported by three motor proteins: kinesin-1, kinesin-2, and dynein; and fewer kinesin-1 motors (1-3 motors per phagosome) are present compared to kinesin-2 (2-6 motors) and dynein (3-10 motors) (13). Yet, MAP7 regulation is primarily mediated through controlling kinesin-1 activity. Mathematical modelling also indicates that regulation is mediated primarily through kinesin-1, as we applied the increase in the binding rate we observed for single kinesin-1 motors to the mathematical model while assuming that kinesin-2 and dynein are unaffected, and the resulting trajectories compare well with our experimental observations (Fig. 4). Tau uses an analogous mechanism to regulate bidirectional motility by inhibiting kinesin-1 processivity while having little direct effect on kinesin-2 and dynein (13). Scaffolding proteins like Miro/Milton and huntingtin that mediate interactions between motor proteins and cargoes have also been proposed to control kinesin-1 activity (38, 39). Together, these results suggest that (1) the net direction of transport of cargoes driven by multiple types of motor proteins can be regulated by controlling the activity of a single motor type, and (2) kinesin-1 activity is a common target for regulation.

Different MAPs have distinct effects on transport. In contrast to MAP7, tau inhibits kinesin-1 processivity to direct bidirectional cargoes towards the microtubule minus end. Tau and MAP7 have distinct localizations in the axon. Tau localizes throughout the axon, but is enriched in the distal axon (40). MAP7 localizes to axonal branches, and contributes to branch formation (18, 19). Interestingly, MAP7 has been shown to compete with tau for binding along the microtubule lattice (23), such that tau and MAP7 may enrich on discrete regions of the microtubule cytoskeleton where they perform distinct functions. For example, tau may act to regulate the balance of plus-and minus-end directed motility along the length of the axon (13). MAP7 may act either to promote movement through the branch site, or potentially guide cargoes into a specific branch. Other MAPs are also likely involved. In the proximal axon and dendrites, MAP2 inhibits kinesin-1 (41) and dynein (42), but enhances the activity of kinesin-3 (43). Collectively, these results suggest that the regulation of organelle transport by MAPs is targeted to both the specific region of the cell, dictated by the localization of MAPs, and to specific motor proteins.

### Experimental procedures

#### Cell culture and phagosome purification

J774A.1 cells (ATCC) were plated in four 100 mm cell culture dishes and were grown in a 37°C incubator at 5% CO_2_ to 80% confluency in DMEM media (Thermo Fisher Scientific), supplemented with 10% FBS (Thermo Fisher Scientific) and 1% Glutamax (Thermo Fisher Scientific). Fluorescent latex beads (500 nm diameter, blue fluorescent, ThermoFisher Scientific) were coated with BSA and incubated with the cells (44). After a 90-min chase, cells were washed with PBS, collected by a cell scraper and washed again with cold PBS, where after each wash the cells were centrifuged at 2000 RPM in 4°C for 5 min and resuspended in PBS. After the second wash, cells were resuspended in motility assay buffer (MAB; 10 mM Pipes, 50 mM K-acetate, 4 mM MgCl_2_, 1 mM EGTA, pH 7.0) supplemented with protease inhibitors, 1 mM DTT, 1 mM ATP, and 8.5% (w/v) sucrose solution in MAB and lysed with a Dounce homogenizer. Following the lysis step, cell lysate was centrifuged at 2000 RPM in 4°C for 10 min and the supernatant (homogenate) was mixed with an equal volume of 62% sucrose. Sucrose solutions were supplemented with ATP and protease inhibitors and loaded in ultracentrifuge tubes (16 x 102 mm – Beckman no. 344061) in the following order: 62% (3.61 mL), homogenate (1 mL), 35% (2.91 mL), 25% (2.91 mL), and 10% sucrose (2.91 mL). The sucrose gradient was centrifuged in a swinging bucket rotor (SW32.1 – Beckman) at 24000 RPM for 72 minutes. After the spin, the fraction containing phagosomes appears as a thin band at the interface between between 25% and 10% sucrose.

#### Protein expression and purification

##### MAP7 expression and purification

MAP7 (accession number BC052637) was cloned from pEGFP-MAP7 (Addgene plasmid #46076) into the baculovirus expression vector pFastBac with N-terminal FLAG and SNAP-tags. Recombinant baculovirus was prepared by standard protocols. *Sf9* cells were infected with recombinant baculovirus for ∼72 hours at 27°C, harvested by centrifugation and re-suspended in lysis buffer (10 mM sodium phosphate, pH 7.5, 0.3 M NaCl, 7% sucrose, 0.5% glycerol, 2 mM DTT, 5 *µ*g/mL leupeptin, 10 mM AEBSF, 10 mM PMSF, and 10 mM TLCK). Cells were lysed by sonication, and centrifuged at 257,000 x g for 35 min. The clarified lysate was added to 4 mL FLAG affinity resin (Sigma-Aldrich) and incubated with rotation at 4°C for 40 min. The resin was transferred to a column and washed with 200 mL FLAG wash buffer (10 mM sodium phosphate, pH 7.5, 0.3 M NaCl, 0.5 mM DTT) and eluted with the same buffer containing 0.1 mg/mL FLAG peptide. Peak fractions were concentrated using an Amicon Ultra-15 centrifugal filter (Millipore) and dialyzed versus 10 mM HEPES, pH 7.5, 0.3 M NaCl, 1 mM DTT, 1 *µ*g/mL leupeptin, 50% glycerol for storage at −20°C.

*Drosophilia melanogaster* ensconsin, isoform A (accession number NP_728941) with an N-terminal 6xHIS tag in pET28a was a gift from Vladimir Gelfand (Northwestern University). E. coli BL21 DE3 cells containing the plasmid were induced with 0.4 mM IPTG and grown at 37°C for 3 hours for protein expression. Cells were pelleted and re-suspended in lysis buffer (10 mM Na phosphate, pH 7.5, 0.4 M NaCl, 0.5% glycerol, 7% sucrose, 7 Mm *β*-mercaptoethanol, 5 *µ*g/mL leupeptin, 10 mM AEBSF, 10 mM PMSF, and 10 mM TLCK) and sonicated. The clarified supernatant was applied to a HIS-Select nickel affinity column (Sigma-Aldrich), and washed with 10 mM Na phosphate, pH 7.5, 0.4 M NaCl, 5 mM imidazole. Bound protein was eluted with the same buffer containing 0.2 M imidazole. Fractions were concentrated by dialysis against 50% glycerol, 10 mM imidazole pH 7.5, 0.4 M NaCl, 1 mM DTT and 1 µg/mL leupeptin and stored at −20°C.

##### Kinesin-1 expression and purification

*Drosophilia melanogaster* or mouse kinesin were cloned into the baculovirus transfer vector pAcSG2 (BD Biosciences). The mouse kinesin heavy chain (accession number BC090841) has a C-terminal biotin-tag followed by a hexa-HIS tag. Its corresponding light chain (accession number BC014845) was cloned with a C-terminal YFP tag. The *Drosophilia* kinesin heavy chain (accession number AF053733) was cloned with a C-terminal biotin-tag followed by a FLAG tag. Its light chain (accession number AF055298) was cloned with a hexa-HIS tag at the C-terminus. The biotin tag is an 88 amino acid sequence segment from the E. coli biotin carboxyl carrier protein, which is biotinylated at a single lysine during expression in *Sf9* cells (45). This tag was used for attachment to streptavidin-conjugated Quantum dots (Qdots). Recombinant baculovirus was prepared by standard protocols.

*Sf9* cells were co-infected with recombinant baculovirus coding for the kinesin heavy chain and light chain, and grown in suspension for ∼72 hours. The mouse kinesin (hexa-HIS tag) was purified by sonicating the infected Sf9 cells in buffer containing 10 mM sodium phosphate, pH 7.5, 0.3 M NaCl, 0.5% glycerol, 7% sucrose, 7 mM *β*-mercaptoethanol, 0.5 mM AEBSF, 0.5 mM TLCK, 5 *µ*g/mL leupeptin, and 1 mM ATP. The cell lysate was clarified at 200,000 x g for 30 min, and the supernatant applied to a HIS-Select nickel affinity column (Sigma-Aldrich). The resin was first washed with buffer A (10 mM sodium phosphate, 10 mM imidazole, pH 7.5, 0.3 M NaCl), and then with buffer A containing 30 mM imidazole. Kinesin was eluted from the column with buffer A containing 200 mM imidazole.

*Drosophilia* kinesin was purified via its FLAG tag. Cells were sonicated in buffer containing 10 mM imidazole, pH 7.0, 0.3 M NaCl, 1 mM EGTA, 5 mM MgCl2, 7% sucrose, 2 mM DTT, 0.5 mM AEBSF, 0.5 mM TLCK, 5 *µ*g/mL leupeptin, and 1 mM ATP. The cell lysate was clarified at 200,000 x g for 30 min, and the supernatant applied to a FLAG affinity resin column (Sigma-Aldrich). The resin was washed with buffer containing 10 mM imidazole, pH 7.0, 0.3 M NaCl, and 1 mM EGTA. Kinesin was eluted from the column by using a 0.1 mg/mL solution of FLAG peptide in wash buffer.

For each purification, the fractions of interest were combined and 1 mM DTT, 1 *µ*g/mL leupeptin, and 10 *µ*M ATP was added prior to concentrating with an Amicon centrifugal filter device (Millipore), Kinesin was dialyzed versus 10 mM imidizole, pH 7.4, 200 mM NaCl, 55% glycerol, 1 mM DTT, 10 *µ*M MgATP and 1 *µ*g/mL leupeptin for storage at −80°C in small aliquots.

##### *In vitro* reconstitution of phagosomes

Bright microtubule seeds were prepared by mixing 25% alexa 647 labeled tubulin and 75% unlabeled tubulin in BRB80 (80 mM K-PIPES, 1 mm MgCl2, 1mM EGTA, pH 6.8) to a final concentration of 5 mg/mL supplemented with 1 mM GTP (Sigma Aldrich) and polymerized at 37°C for 20 min. After polymerization, bright microtubule seeds were dissolved in cold BRB80 supplemented with 1 mM GMPCPP (Jena Bioscience) and 2 mM MgCl2. This mixture was incubated at 37°C for 30 min and then pelleted at 100,000 x g for 10 min at 4°C in TLA100 rotor (Beckman). The pellet was resuspended in 70 *µ*L cold BRB80 and incubated on ice for 20 minutes for depolymerization. The solution was centrifuged at 100,000 x g for 10 min at 4°C and resuspended with 2 mM MgCl2 and 1 mM GMPCPP. The pellet was then resuspended in BRB80 and aliquoted in 5 µL stocks and flash frozen. To prepare polarity marked microtubules, GMPCPP seeds were warmed at 37°C for 1 min to which 8% labeled tubulin was added in 92% unlabeled tubulin in BRB80 and 1 mM GTP to a final concentration of 2 mg/mL. After 25 min incubation, these microtubules were stabilized with 20 *µ*M taxol (Cytoskeleton) and incubated for another 25 min at 37°C. Silanized coverslips were mounted on glass slides using vacuum grease and double sided tape to form 20-25 *µ*L flow chambers.

After flowing 1 chamber volume of anti-*β*-tubulin (2.5:50 in BRB80), the chamber was surface treated with F-127 (Sigma) to block any non-specific binding. Diluted microtubules (2:50 in BRB80 supplemented with Taxol) were flown through the chamber, following a wash with two chamber volumes of BRB80 supplemented with 20 µM taxol. Purified phagosomes were added to the chamber, supplemented with 0.2 mg/mL BSA, 10 mM DTT, 1 mm MgATP, 20 µM taxol, 15 mg/mL glucose, >2000 units/g glucose oxidase, >6 units/g catalase, and 1 mg/mL casein. For MAP7 experiments, microtubules were mixed with 10 nM MAP7 and incubated in the flow chamber for 15-20 minutes.

##### Kinesin-1/bead complex preparation

Anti-GFP coated carboxylated polystyrene beads (500 nm diameter) were prepared as described (46). 3.9 mM full length kinesin1–YFP was titrated to 3 different concentrations – 200 nM, 400 nM, and 800 nM, and incubated with 7.28 x 10^8^ beads. This translated the motor/bead ratio to 5000 kinesin-1/beads (for 200 nM), 10,000 kinesin-1 beads (for 400 nM), and 15,000 kinesin-1/beads (for 800 nM). 0.25 *µ*M ATP, 0.4 mg/mL BSA, and 2.5 *µ*L of anti-GFP coated beads were mixed and the solution was then sonicated for 1 minute. Afterwards, full length kinesin–1 (with above stated concentrations) was added to the solution and gently mixed (abrasive pipetting may damage the anti–GFP coated beads). The final solution was then incubated for 20 minutes in 4°C. Full length Kinesin–1 coated beads were added to the chamber with immobilized microtubules supplemented with 0.4 mg/mL BSA, 1 mM MgATP, 10 µM taxol, 15 mg/mL glucose, *>*2000 units/g glucose oxidase, and *>*5 units/g catalase.

##### Optical Trapping

The optical trap was built on an inverted microscope (Eclipse Ti-E: Nikon) with a 1.49 NA oil-immersion objective. A 1064 nm laser beam (IPG Photonics) was expanded to overfill the back aperture of the objective. To measure the bead displacement and force, a quadrant photodiode was positioned conjugate to the back focal plane of the condenser. The optical trap stiffness (*K*_*T* *RAP*_) and position calibration (*β*) were determined by fitting a Lorenzian function to the power spectrum of the bead’s thermal fluctuations. Force events are defined as displacement from the trap center where the force is greater than 0.5 pN. Short and low-force events are common, due to the detachment of motor teams befor they their maximal force. To this end, we analyzed optical trap data at low (*T*_*ST* *ALL*_ *≥* 70 msec), intermediate (*T*_*ST* *ALL*_ *≥* 300 msec), and high (*T*_*ST* *ALL*_ *≥* 1000 msec) stall times. Low stall times indicate cases where teams of motors are engaged under load but rarely reach stall. On the other hand, a high stall time indicates cases where teams of motors are likely exerting their maximal forces (stall force). In the case of intermediate stall times, motors are expected to exert detachment and stall forces. Since our force data indicate a subpopulation of detachment and stall forces, we used a probabilistic model (Bayesian Gaussian Mixture Model) to analyze the expected values from the force histogram and used bootstrapping to determine the optimal number of components in our data set.

##### Single molecule kinesin-1 motility assay

The procedure was essentially as described previously (47). Briefly, 1 *µ*L of 0.2 *µ*M streptavidin Quantum dot (Thermo Fisher Scientific, Waltham, MA) was mixed with 1 *µ*L of 0.2 *µ*M *Drosophilia* kinesin (with C-terminal biotin tag) for 10 min at room temperature. The mixture was then diluted 200-400 fold in 80 mM PIPES, pH 6.9, 1 mM MgCl2, 1 mM EGTA, 1 mM MgATP, and an oxygen scavenging system composed of 0.1 mg/mL glucose oxidase, 0.02–0.18 mg/mL catalase and 3 mg/mL glucose) and flowed into a flow cell with microtubules attached to the surface via N-ethyl maleimide-modified myosin. The slide was mounted onto an inverted Nikon TE2000U Microscope equipped with a Nikon objective lens (100X; numerical aperture, 1.49 NA) for through-the-objective total-internal-reflection fluorescence (TIRF) microscopy. Single molecule assays were performed at room temperature (25 ± 1°C). Quantum dots and microtubules were excited with a 488nm argon ion Spectra Physics laser. Fluorescence images of Quantum dots and microtubules were then split by color and projected side by side onto a Stanford Photonics Mega 10X camera. For typical experiments, 500-1000 images were captured at 10 frames/sec with an exposure time of 100ms. Using 2×2 binning, the pixel resolution is 117.6 nm/pixel. Images were then imported into ImageJ (National Institutes of Health, Bethesda, MD) for processing. Those events in which the kinesin moves on microtubules in both directions for at least 300nm distance are classified as diffusive events. Those events in which kinesin only moves in one direction were classified as processive events. Run lengths were determined manually. Similar procedures were used to image full-length mammalian kinesin-1-YFP. In these experiments, exposure times were increased to 1s.

##### Multiple-motor based motility assay

Mammalian kinesin-1 coated beads with a titration of 10,000 kinesin-1/beads held with a low stiffness optical trap were brought down to the microtubules and allowed to run out of the trap (12). Their motility was analyzed using FIESTA (Fluorescence Image Evaluation Software for Tracking and Analysis) (48). Position vs time trajectories from FIESTA tracking were compared to kymographs to validate the automated tracking. Run lengths were determined as net displacements from the trap center to the maximum position before the bead either detaches from the microtubule or slips back to the center of the trap. The 1-dimensional mean squared displacement (MSD = 2Dt^*α*^) was calculated using internal averaging, where D is the diffusion coefficient, t is the time interval, and *α* is the scaling exponent corresponding to stationary (*α*=0), diffusive (*α*=1), and processive (*α*=2) movement (49).

##### Statistical analysis

All the data is presented as the mean plus/minus standard error of mean (SEM). The number of experiments and recordings (*n*) are mentioned in the figure legends. Statistical analysis was performed using MATLAB (Mathworks Inc.) Statistical analysis was performed using One-Factor ANOVA, student’s t-test, bootstrapping and Efron’s percentile method, Kolmogorov-Smirnov test, and Bayesian Information Criterion. Key findings from this study are summarized in the Supplemental information (Table S1 and S2).

## Acknowledgements

The authors thank Gary Brouhard (McGill University, Montreal, Canada) for providing reagents, Christophe Leterrier (NeuroCyto, INP CNRS-Aix Marseille Université, Marseille, France) for providing the image of a neuron (Fig. 4F), Ricardo Henriques for providing the LaTeX template, and Linda Balabanian for preparing anti-GFP coated carboxylated polysterene beads. We are also grateful to Malina Iwanski and Loïc Chaubet for providing valuable input on the manuscript. AGH is supported by the Natural Sciences and Engineering Research Council of Canada (RGPIN-2014-06380) and the Canadian Institutes of Health Research (PJT159490); KMT is supported by NIH (GM078097).

## Author contributions

A.R.C., K.M.T, and A.G.H. designed research. A.R.C. performed optical trapping assays. H.L. performed single molecule kinesin-1 motility assays. E.B.K. and C.S.B. expressed and purified kinesin-1 and MAP7. A.R.C. performed phagosome purifications. A.R.C., A.G.H., H.L., and K.M.T. analyzed the data and wrote the manuscript. A.G.H and K.M.T. conceptualized the project and supervised the work.

## Supporting Information

**Movie S1: Motility of kinesin–1 with 0 nM MAP7**

A single full length mammalian kinesin-1 (red) is moving along the microtubule lattice (green) with no MAP7 (Scale bar = 3*µ*m).

**Movie S2: Motility of kinesin–1 with 10 nM MAP7**

The frequency of kinesin motors (red) moving along the microtubule lattice (green) increases in the presence of 10 nM MAP7 (not shown) (Scale bar = 3*µ*m).

**Movie S3: Motility of kinesin–1 with 20 nM MAP7**

Compared to motility of kinesin-1 with 10 nM MAP7, a higher fraction of single kinesins (red) can be seen moving diffusively along the microtubules (green) (Scale bar = 3*µ*m).

**Movie S4: Motility of teams of FL Kinesin–1 with 0 nM MAP7**

Beads coated with full length mammalian kinesin-1 motors that exhibit motility by ∼1-2 kinesin-1 motors were held with a low stiffness optical trap and brought down to the microtubules (not shown) and allowed to run out of the trap. In this case no MAP7 was added to the microtubules. A high frequency of detachment and reattachment events by kinesin motors can be observed.

**Movie S5: Motility of teams of FL Kinesin–1 with 10 nM MAP7**

Compared to the motility of teams of kinesin-1 without MAP7, we can observe that kinesins remain engaged to the microtubule for longer periods of time and do not detach.

**Movie S6: MAP7 imaging on microtubules**

MAP7 forms static patches along the microtubule lattice (not shown).

**Table S1: Summary of optical trap data**

**Table S2: Summary of motility data**

### 1. MAP7 imaging and dwell time analysis

MAP7 imaging was performed in TIRF using a 647 nm laser with an exposure time of 500 msec for 5 mins. The trajectories of bound MAP7 were analyzed from kymographs **(Fig. S1J)**. Interestingly, we observed that MAP7 remains bound to the microtubules longer than our imaging times. To take into account the dwell time of microtubule bound MAP7 prior or subsequent to the imaging, we used survival analysis. Assuming that every MAP7 event follows the same survival function, *S*(*t*), we estimated the dwell time distribution using the Kaplan-Meier estimator [4]:

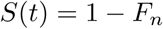

Here *F*_*n*_ is the empirical cumulative distribution of the observable MAP7 dwell time. To measure the mean dwell time, we fit the Weibull survivor function to the distribution, *S*(*t*). Briefly, the weibull distribution is defined as follows:

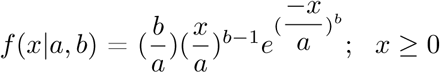

Here, *a* and *b* are the scale and shape parameters, respectively. The mean and standard deviation of the weibull distribution with parameters, a and b, were computed via MATLAB (Mathworks Inc.) as follows:

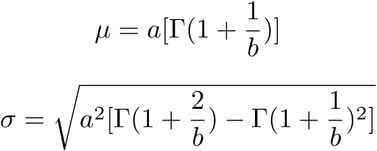

Here, Γ is the Gamma function.

### 2. Relative Binding rate

Binding rate is defined as the time it takes to change the state of an unbound diffusive motor to a bound processive motor (**Fig. 1F**). In a stationary optical trap, binding rate, π (*s*^-1^), can be determined as:

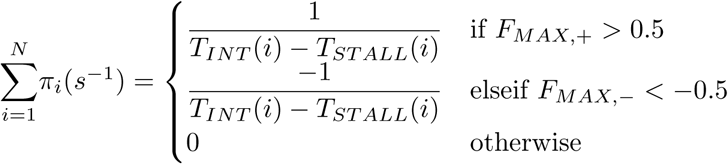

*N* = Total number of force events in an optical trap recording

*T*_*ST* *ALL*_ = Duration of a force event

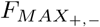= Maximum Force in the plus or minus-end

*T*_*INT*_ = Time interval of the *i*^*th*^ loop

### 3. Step Size analysis

As cargoes move processively, molecular motors take discrete steps along the microtubules. To determine the stepping behavior of molecular motors, we computed the pairwise distribution using the distance vs. time data of an optical trap trajectory **(Fig. S1H)**. We computed the step size by pairwise distribution since the positional noise of the cargo is much noisier than the step function. To do so, we thresholded force traces, 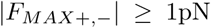, and smoothed the position data using a Savitsky Golay filter. To determine the pairwise distance for a single trace, we take the difference between the first point and the point one time step after it. The difference in position between pairs of points is binned and the distribution has multiple peaks. The maximum peak refers to most frequent step size and is recorded as the average step size.

### 4. Force-Velocity analysis

Normal smoothing averaging techniques such as Savitsky-Golay filtering assign equal weights to each point falling in a specific window size that is defined by the user. Although this assumption might work for input data where noise does not change over time, force data that outputs from the quadrant photodiode has variable noise over time. To take into account some central points that might be noisier than some other ones, we filtered force traces at a bandwidth of 40 Hz via the Nadaraya-Watson kernel regression filter [6]:

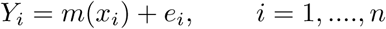

Where, *e*_*i*_ denotes the errors that satisfy a normal distribution, *n* denotes the number of points over which the regression is computed (*n* = 100), and *m*(*x*) is a weight function given by:

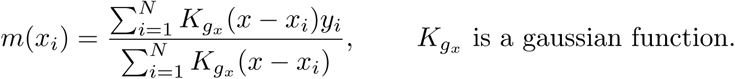

To compute the velocity, we di?erentiated position data through a discrete wavelet transform (DWT) with Biorthogonal spline wavelet (B-spline).

---

**Figure S1:**
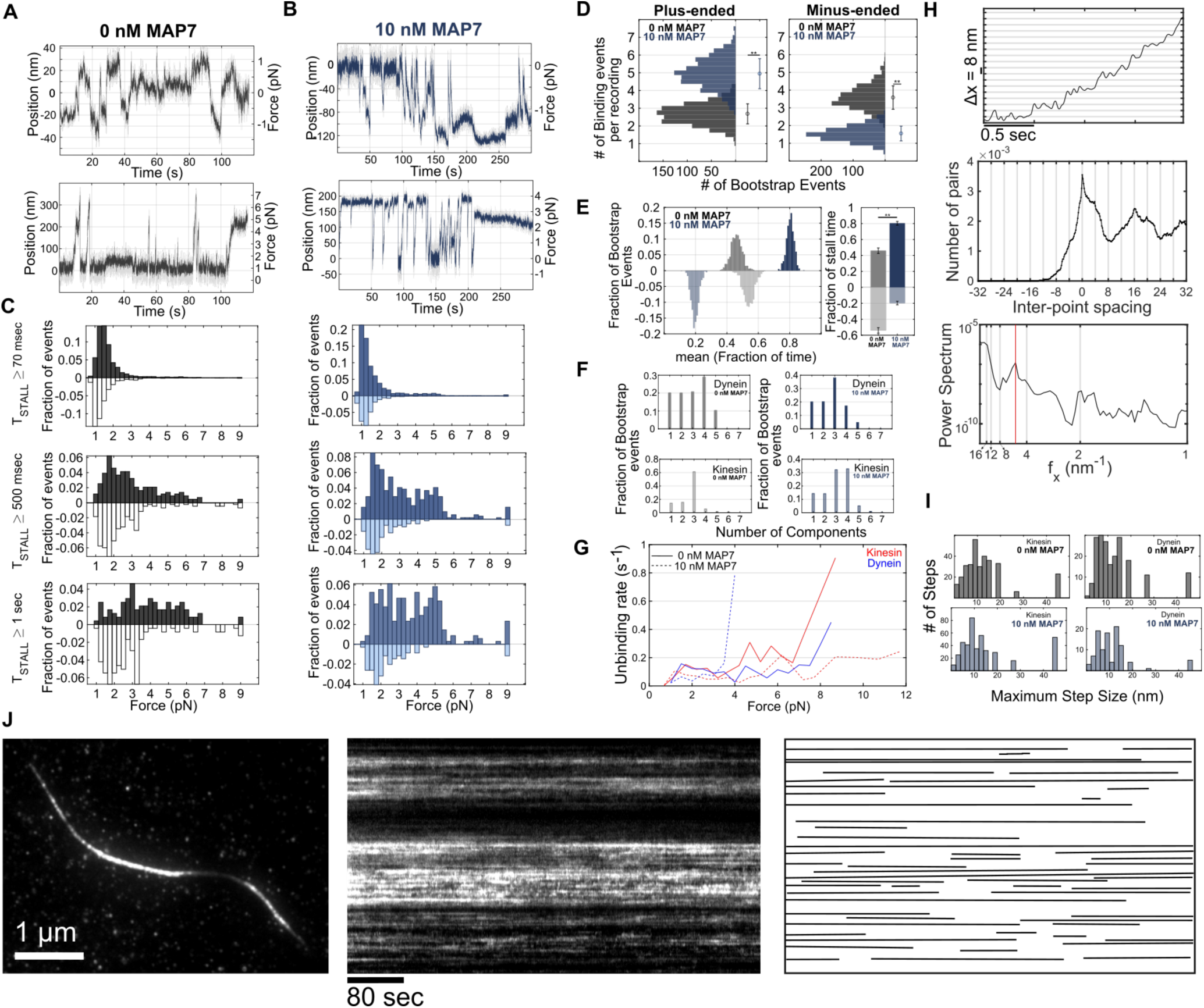
MAP7 biases cargo motility towards the MT plus-end. **(A,B)** In addition to long and stall-force events, short and low-force events are also common due to detachment before motor teams reach their maximal stall force. **(C)** Force events are defined as displacements from the trap center where the force is > 0.5 pN. Kinesin and dynein forces are approximately balanced without MAP7. With MAP7, kinesin forces increase in frequency, while dynein-mediated forces decrease in frequency and magnitude. This effect can be observed for events with short and long stall times. **(D)** Bootstrap analysis of the number of binding events per recording shows that, on average, kinesin binds to the microtubule more frequently and dynein binds less frequently in the presence of MAP7. **(E)** Bootstrap analysis of the fraction of time of force events. Our results indicate that MAP7 increases the time plus-ended motors remain active on microtubules by ∼25%. **(F)** Bootstrap analysis of bayesian information criterion for determining the optimal number of components to describe the force histograms (**Fig. 1C**). **(G)** Motor unbinding rate was determined as described [1]. Our results indicate that MAP7 does not a?ect kinesin or dynein unbinding rate at zero load (ϵ_[0*nM,*+]_ = 0.0037*/s,* ϵ_[0*nM,-*]_ = 0.0237*/s,* ϵ_[10*nM,*+]_ = 0.0058*/s,* ϵ_[10*nM,-*]_ = 0.0207*/s*). **(H and I)** Step size distribution via pairwise distance calculation shows that motor stepping behavior is unaffected by MAP7. **(J)** MAP7 binds statically along the microtubule lattice.

**Figure S2:**
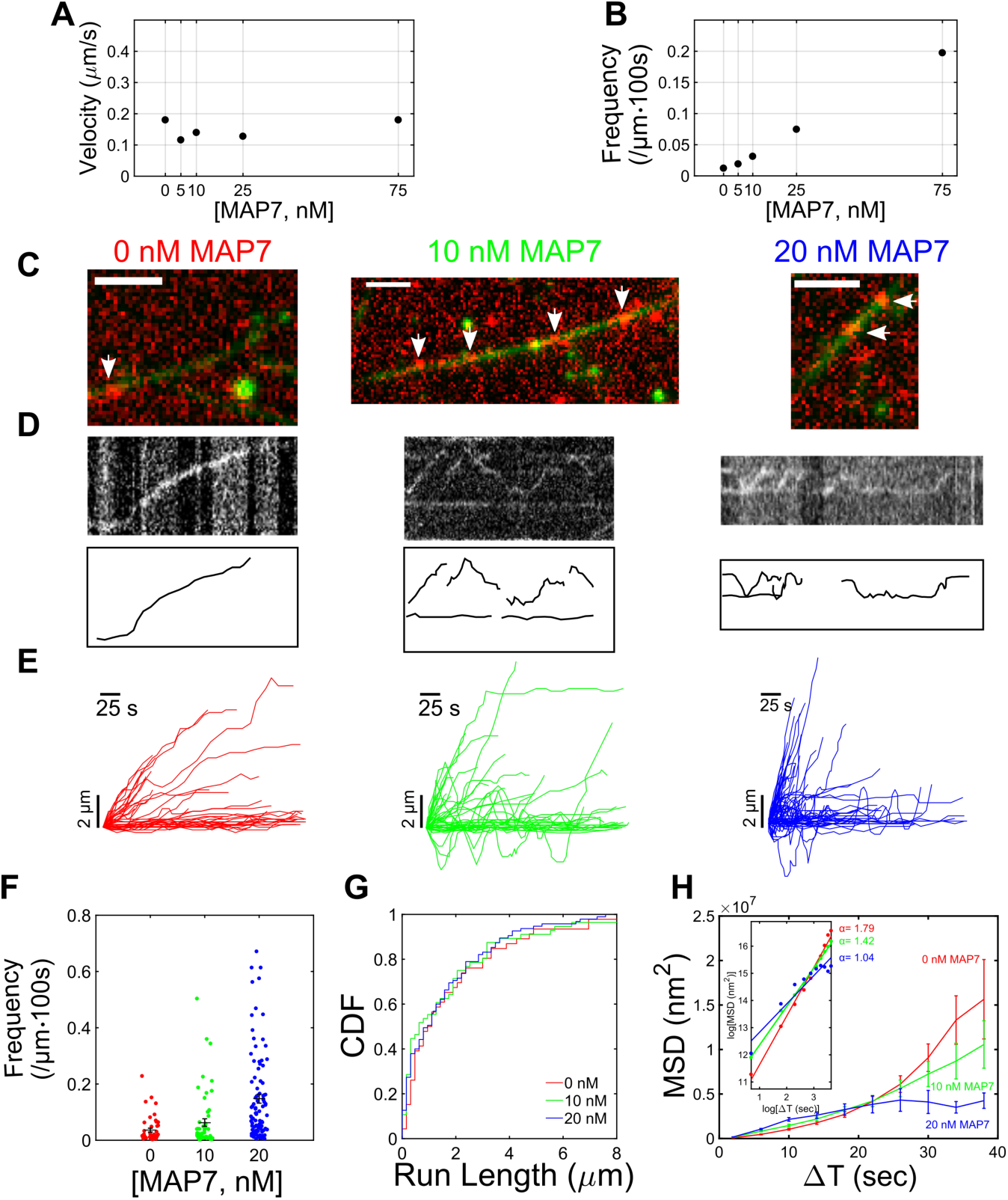
MAP7 increases the engagement time of kinesin-1 motors on microtubules. **(A)** *In vitro* motility assays indicate that MAP7 does not influence the velocity of single molecules of kinesin-1. **(B)** On the other hand, MAP7 does influence the binding of kinesin-1 by increasing the frequency of both di?usive and processive runs. To validate whether mammalian full length kinesin-1 behaves similarly to *Drosophilia* full length kinesin-1, we conducted motility assays with mammalian full length kinesin-1 constructs. (**C,D,E**) Motility assays and kymographs indicate that MAP7 increases the processive and diffusive activity of kinesin-1 motors on the microtubules. (**F**) We further assessed the number of events per *µ*m per 100 sec and observe that, similar to *Drosophilia* kinesin-1, mammalian kinesin-1 also binds more frequently to the microtubule. (**G**) Kinesin-1 run lengths remain unchanged in the presence of MAP7. It should be noted that these run lengths are a mix of diffusive and processive runs (0 nM MAP7= 1848 +/-297 nm, 10 nM MAP7 = 1639 +/-286 nm nm, 20 nM MAP7 = 1587 +/-179.19 nm). (**H**) MSD analysis indicates that mammalian kinesin-1 becomes more diffusive in the presence of MAP7. (0 nM MAP7: *n* = 46 runs from 26 microtubules; 10 nM MAP7: *n* = 56 runs from 23 microtubules; 20 nM MAP7: *n* = 95 runs from 31 microtubules). Scale bar = 3*µ*m.

**Figure S3:**
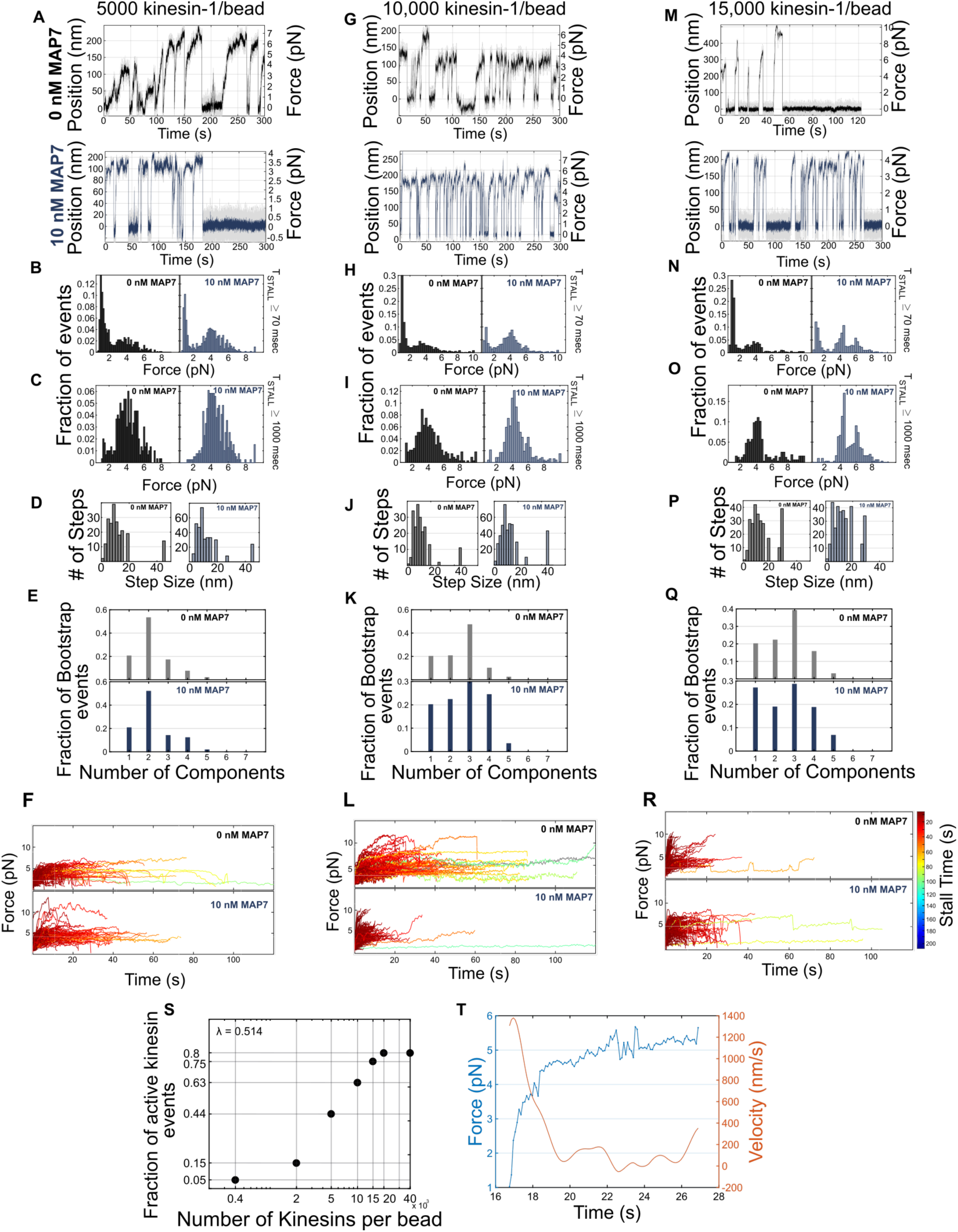
MAP7 increases the binding rate of single and teams of kinesin-1 motors to the microtubules. **(A,G,M)** Example force traces by single and teams of full length kinesin-1 motors. Single **(B,C)** and teams of kinesin-1 motors **(H,I,N,O)** exert short and low-force events along with long and stall force events. In the presence of MAP7, these short and low-force events are diminished. **(D,J,P)** MAP7 does not affect the stepping behavior of kinesin-1 motors. **(E,K,Q)** Bootstrap analysis of bayesian information criterion was used for determining the optimal number of components to describe the force histograms (**Fig. 3C,I,O**). **(F,L,R)** Trajectories of single and teams of kinesin-1 motor force events. Color bar indicates the engagement time of kinesin motors on the microtubule. (**S**) We varied the concentration of kinesins per bead to analyze the fraction of active beads. The motile fraction of beads driven by at least 1 kinesin can be modeled by a Poisson CDF: *P* (≥ 1) = 1-*e*^*-λx*^ [2, 5]. In our system, we expect to observe at least one active kinesin per bead 50% of the time. **(T)** Example of a force-velocity event by a single kinesin reaching stall force under a static optical trap. As motors reach stall or detachment force, they slow down, indicating an inverse relationship between load and velocity.

**Figure S4:**
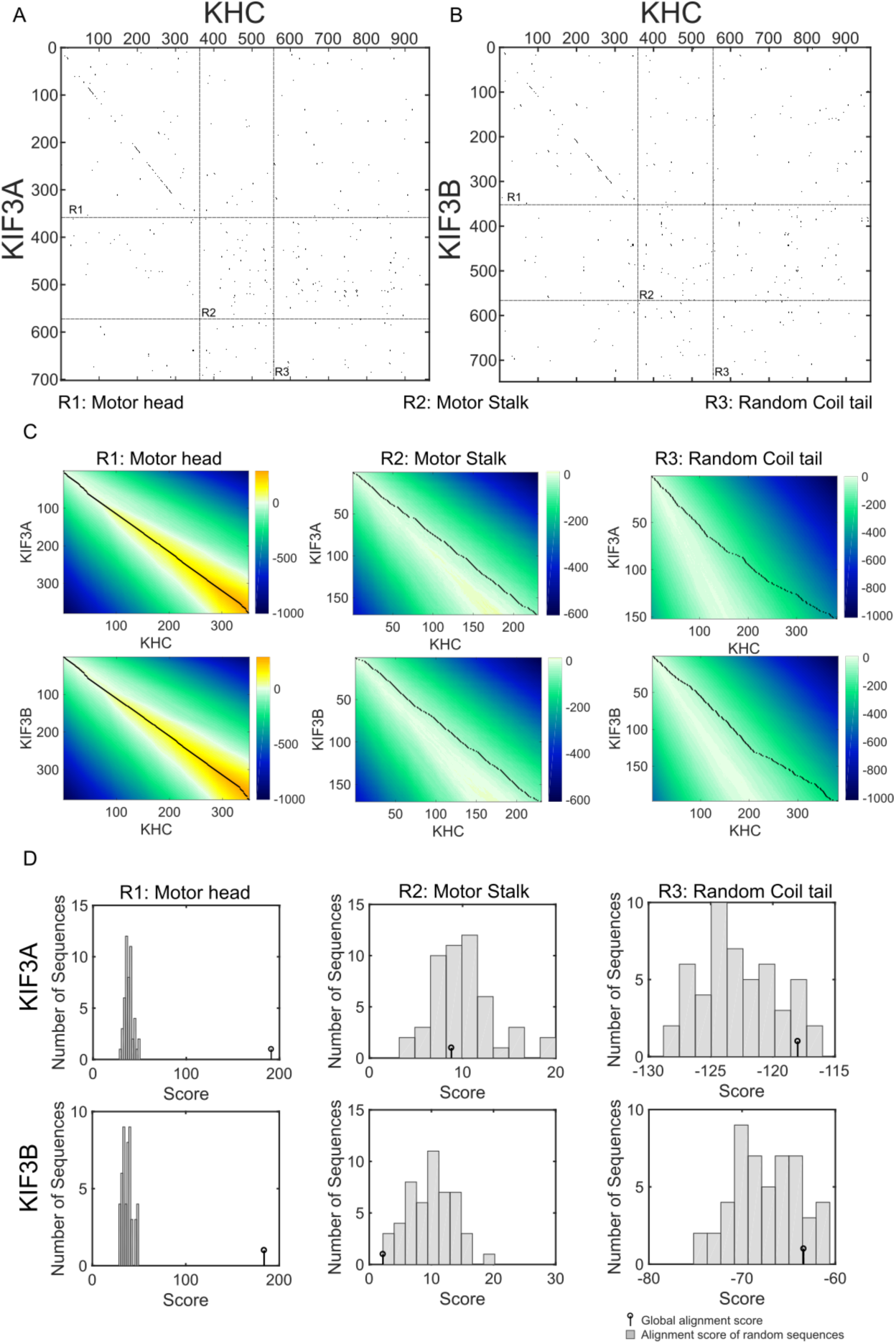
Kinesin-2 is unaffected by MAP7. (**A,B**) Using the Needleman-Wunsch algorithm [3], we aligned the Kinesin heavy chain (KHC) with KIF3A and KIF3B. The stalk domain of kinesin-1 has been shown to interact with MAP7 [7]. (**C**) Our results show that the motor head domain of KIF3A/B align well with KHC, while the stalk and random coil tail domain sequence of KIF3A/B and KHC have no homology. The color gradient here indicates the score space, while the dashed line shows the winning path. As seen from this figure, the score space aligns well with the winning path for KIF3A/B motor head (indicating high homology between KHC and KIF3A/B motor head), while the score space and winning path do not align well between KIF3A/B and KHC stalk and random coil tail domain. (**D**) To assess the statistical significance between the homology and non-homology sequence and validate that homology is not due to randomness of sequence, we shuffled sequences of KIF3A/B motor head, stalk, and random coil tail domains. Our results suggest that the homology and non-homology between KIF3A/B and KHC is not due to random sequences as demonstrated by the difference between actual score (stem) and the monte-carlo simulated score from randomized sequence (histogram).

